# Assessing genomic diversity and signatures of selection in Original Braunvieh cattle using whole-genome sequencing data

**DOI:** 10.1101/703439

**Authors:** Meenu Bhati, Naveen Kumar Kadri, Danang Crysnanto, Hubert Pausch

**Affiliations:** Animal Genomics, ETH Zürich, Zürich, Switzerland

**Author notes:** Authors and email addresses: Meenu Bhati, Naveen Kumar Kadri, Danang Crysnanto, Hubert Pausch.

## Abstract

**Background:** Autochthonous cattle breeds represent an important source of genetic variation because they might carry alleles that enable them to adapt to local environment and food conditions. Original Braunvieh (OB) is a local cattle breed of Switzerland used for beef and milk production in alpine areas. Using whole-genome sequencing (WGS) data of 49 key ancestors, we characterize genomic diversity, genomic inbreeding, and signatures of selection in Swiss OB cattle at nucleotide resolution.

**Results:** We annotated 15,722,811 million SNPs and 1,580,878 million Indels including 10,738 and 2,763 missense deleterious and high impact variants, respectively, that were discovered in 49 OB key ancestors. Six Mendelian trait-associated variants that were previously detected in breeds other than OB, segregated in the sequenced key ancestors including variants causal for recessive xanthinuria and albinism. The average nucleotide diversity (1.6 × 10^-3^) was higher in OB than many mainstream European cattle breeds. Accordingly, the average genomic inbreeding quantified using runs of homozygosity (ROH) was relatively low (F_ROH_=0.14) in the 49 OB key ancestor animals. However, genomic inbreeding was higher in more recent generations of OB cattle (F_ROH_=0.16) due to a higher number of long (> 1 Mb) runs of homozygosity. Using two complementary approaches, composite likelihood ratio test and integrated haplotype score, we identified 95 and 162 genomic regions encompassing 136 and 157 protein-coding genes, respectively, that showed evidence (*P* < 0.005) of past and ongoing selection. These selection signals were enriched for quantitative trait loci related to beef traits including meat quality, feed efficiency and body weight and pathways related to blood coagulation, nervous and sensory stimulus.

**Conclusions:** We provide a comprehensive overview of sequence variation in Swiss OB cattle genomes. With WGS data, we observe higher genomic diversity and less inbreeding in OB than many European mainstream cattle breeds. Footprints of selection were detected in genomic regions that are possibly relevant for meat quality and adaptation to local environmental conditions. Considering that the population size is low and genomic inbreeding increased in the past generations, the implementation and adoption of optimal mating strategies seems warranted to maintain genetic diversity in the Swiss OB cattle population.

## Introduction

Following the domestication of cattle, both natural and artificial selection led to the formation of breeds with distinct phenotypic characteristics including morphological, physiological and adaptability traits [1]. With an increasing demand for animal-based food products, few breeds were intensively selected for high milk (e.g., Holstein, Brown Swiss) and beef (e.g., Angus) production. The predominant selection of cattle from specialized breeds caused a sharp decline in the population size of local cattle breeds [2, 3]. Although less productive under intensive production conditions, local breeds of cattle might carry alleles that enable them to adapt to local conditions. Therefore, local breeds represent an important genetic resource to facilitate animal breeding in the future under challenging and changing production conditions [4, 5]. Characterizing the genetic diversity of local cattle breeds is important to optimally manage these genetic resources.

The Swiss Original Braunvieh (OB) cattle breed is a dual purpose taurine cattle breed that is used for beef and milk production in alpine areas [6, 7]. In transhumance farming, the cattle graze at alpine pastures (between 1000 and 2400 meter above sea level) during the summer months and return to the stables for the winter months [7]. Mainly due to their strong and firm legs and claws, they are well adapted to the alpine terrain. Under extensive farming conditions, OB cattle may outperform specialized dairy breeds in terms of fertility, longevity and health status [8]. However, in the early 1960s, Swiss cattle breeders began inseminating OB cows with semen from US Brown Swiss sires to increase milk yield, reduce calving difficulties and improve mammary gland morphology of the Swiss OB cattle population [9]. The extensive cross-breeding of OB cows with Brown Swiss sires decreased the number of female OB calves entering the herdbook to less than 2,000 by mid 1990’s [9] (Additional file 1). Since then, the size of the Swiss OB population increased steadily, facilitated by governmental subsidies.

A number of studies investigated the genomic diversity and population structure of the Swiss OB cattle breed using either pedigree or microarray data [9, 10]. In spite of the small population size, genetic diversity is higher in OB than many commercial breeds likely due to the use of many sires in natural mating and lower proportion of artificial insemination [9, 10]. Genomic inbreeding and footprints of selection have been compared between OB and other Swiss cattle breeds using SNP microarray-derived genotypes [10]. Because the SNP microarrays were designed in a way that they interrogate genetic markers that are common in the mainstream breeds of cattle, they might be less informative for breeds of cattle that are diverged from the mainstream breeds [11]. Ascertainment bias is inherent in the resulting genotype data because rare, breed-specific, and less-accessible genetic variants are underrepresented among the microarray-derived genotypes [12]. This limitation causes observed allele frequency distributions to deviate from expectations which can distort population genetics estimates [13].

With the availability of whole genome sequencing (WGS), it has become possible to discover sequence variant genotypes at population scale [14]. While sequence variant genotypes might be biased toward the reference allele, this reference bias is less of a concern when the sequencing coverage is high [15]. According to Boitard et al. 2016 [16], WGS data facilitate detecting selection signatures at higher resolution than SNP microarray data. Moreover, the WGS-based detection of runs of homozygosity (ROH) is more sensitive for short ROH that are typically missed using SNP microarray-derived genotypes.

In the present study, we analyze more than 17 million WGS variants of 49 key ancestors of the Swiss OB cattle breed that were sequenced to an average fold-coverage of 12.75 per animal [17]. These data enabled us to assess genomic diversity and detect signatures of past or ongoing selection in the breed at nucleotide resolution. Moreover, we estimate genomic inbreeding in the population using runs of homozygosity.

## Material and Methods

### Sequence variant genotyping

We considered genotypes at 17,303,689 biallelic variants (15,722,811 SNPs and 1,580,878 Indels) that were discovered and genotyped previously [17] in the autosomes of 49 key ancestors of the OB population using a genome graph-based sequence variant genotyping approach [18]. In brief, 49 OB cattle were sequenced at between 6 and 38-fold genome coverage using either Illumina HiSeq 2500 (30 animals) or Illumina HiSeq 4000 (19 animals) instruments. The sequencing reads were filtered and subsequently aligned to the UMD3.1 assembly of the bovine genome [19] using the mem-algorithm of the Burrows Wheeler Aligner (BWA) software package [20]. Single nucleotide and short insertion and deletion polymorphisms were discovered and genotyped using the Graphtyper software [18]. Following recommended filtration criteria (see Crysnanto et al. [17] for more details), 15,722,811 SNPs and 1,580,878 Indels were retained for subsequent analyses. Beagle [21] phasing and imputation was applied to improve the primary genotype calls from Graphtyper and infer missing genotypes. Unless stated otherwise, imputed genotypes were considered for subsequent analyses, because Beagle imputation considerably improved the primary genotype calls particularly in samples that had been sequenced at low coverage [17].

### Variant annotation and evaluation

Functional consequences of 15,722,811 SNPs and 1,580,878 Indels were predicted according to the Ensembl (release 91) annotation of the bovine genome assembly UMD3.1 using the Variant Effect Predictor tool (VEP v.91.3) [22] with default parameter settings. The impacts of amino acid substitutions on protein function were predicted using the sorting intolerant from tolerant (SIFT) (version 5.2.2) [23] algorithm that has been implemented in the VEP tool. Variants with SIFT scores less than 0.05 were considered to be likely deleterious to protein function. In order to assess if known Mendelian trait-associated variants segregate among 49 sequenced OB cattle, we downloaded genomic coordinates of 155 trait-associated variants that are curated in the Online Mendelian Inheritance in Animals (OMIA) database [24, 25].

### Population genetic analysis

Nucleotide diversity (π) quantifies the average number of nucleotide differences per site between two DNA sequences that originated from the studied population [26]. We estimated π of the OB population over the entire autosomal genome using VCFtools v0.1.15 (in windows of 10 kb) [27].

### Detection of runs of homozygosity

Runs of homozygosity were identified using a Hidden Markov Model (HMM)-based approach implemented in the BCFtools/RoH software [28, 29]. The recombination rate was assumed to be constant along the genome at 10^-8^ per base pair (1 cM/Mb). For the HMM-based detection of ROH, we considered phred-scaled likelihoods (PL) and allele frequencies of 15,722,811 filtered SNPs before Beagle imputation. Because samples that are sequenced at low coverage are enriched for ROH [30], we considered only 33 samples with average sequencing coverage greater than 10-fold for the detection of ROH (Additional file 2). We only considered ROH longer than 50 kb because they were less likely to contain false-positives (Phred-score > 67 in our data, Additional file 2). Genomic inbreeding (F_ROH_) was calculated for each animal as F_ROH_=∑L_ROH_/L_GENOME_, where ∑L_ROH_ is the length of all ROH longer than 50 kb and L_GENOME_ is the length of the genome covered by SNPs [31], which is 2,512,054,768 bp in our data.

Further, ROH were classified into short (50 - 100 kb), medium (0.1 - 2 Mb) and long ROH (>2 Mb) reflecting ancient, historical, and recent inbreeding, respectively [32]. The contribution of each ROH category to F_ROH_ was calculated for each animal. Average genomic inbreeding was compared between animals born before and after 1989 using the two samples t-test.

### Detection of signatures of selection

To avoid potential bias arising from extended relationships among the sequenced animals, we did not consider nine sons from sire-son pairs for the detection of signatures of selection. For the remaining 40 cattle, we considered genotypes at 9,051,833 SNPs for which the ancestral allele provided by Rocha et al. [33] was detected in at least two species other than cattle and where it agreed with either the reference or alternate allele in our data. Haplotypes were phased using the Eagle2 software [34] with default parameter settings and assuming a constant recombination rate along the chromosome.

### Integrated haplotype score (iHS)

To identify signatures of ongoing selection, integrated haplotype scores (iHS) were calculated for 8,465,912 variants with minor allele frequency (MAF) greater than 0.01 using the R package rehh v.2 [35]. We obtained iHS that ranged from −6.6 to 6.4. Subsequently, |iHS| were averaged for non-overlapping windows of 40 kb over the whole genome. Windows with either less than 10 SNPs were removed. To test if variants with similar |iHS| properties were pooled in 40 kb windows, we followed the approach of Granka et al. [36]. Specifically, we randomly selected the same number of SNPs that were pooled in 40 kb windows and calculated the mean variance of |iHS| in the true and permuted 40 kb windows for each chromosome. This procedure was repeated for 10,000 randomly selected 40 kb windows. The variance of |iHS| in the non-overlapping 40 kb windows (0.24) was significantly (*P* < 0.01) less than in windows of randomly selected SNPs (0.37) indicating that SNPs that were grouped in 40 kb windows had |iHS| values that were more alike than random SNPs.

### Composite likelihood ratio (CLR)

Composite likelihood ratio (CLR) tests were carried out to identify alleles that are either close to fixation or already reached fixation due to past selection. Following the recommendation of Huber et al. [37], we removed 118,124 SNPs from the data which were fixed for the ancestral alleles because such sites are not informative for CLR tests. Using a pre-computed empirical allele frequency spectrum of 8,933,709 SNPs for which ancestral and derived alleles were assigned (see above), we calculated CLR statistics in non-overlapping 40 kb windows using SweepFinder2 [38, 39]. A window size of 40 kb was chosen to allow comparison and alignment between |iHS| and CLR values.

Empirical *P* values were calculated for CLR and |iHS| windows [40] and the top 0.5% of windows of each statistic were considered as candidate signatures of selection. Adjacent top 0.5% windows were merged separately for each statistic using BEDTools v2.27.1 [41]. For each merged candidate signature of selection, the lowest *P* value among the merged windows was retained.

### Characterization of signatures of selection

Genes within candidate signatures of selection were determined based on the Ensembl (release 91) annotation of the UMD3.1 assembly of the bovine genome. Gene-set enrichment analysis of genes within candidate signatures of selection was performed using PANTHER v.14.1 [42]. Specifically, we investigated if these genes were enriched in the functional categories of GO-slim Biological Process and PANTHER pathways using *P* ≤ 0.05 as significance level.

To determine the overlap between QTL and candidate signatures of selection, we downloaded genomic coordinates for 122,893 QTL from the Animal QTL Database [43, 44]. We classified 85,722 unique QTL that were located on the 29 autosomes into six trait categories: exterior, health, milk, meat and carcass, production and reproduction (Additional file 3). QTL with identical genomic coordinates in associated trait categories were considered as one QTL. We used the intersect module of BEDTools v2.27.1 [41] to identify QTL that overlapped with CLR and |iHS| candidate regions for each of the six trait categories separately. To test if QTL were enriched in candidate signatures of selection, we used a permutation test with 10,000 permutations. In each permutation, we randomly sampled the same number of regions of the same size as the candidate signatures of selection from CLR and |iHS| for each chromosome separately, and overlapped them with QTL of the respective trait categories using BEDTools (see above). The number of QTL that overlapped permuted regions was used as the empirical null distribution to calculate *P* values. *P* values less than 0.05 were considered as indicators for a significant enrichment of QTL in candidate signatures of selection.

## Results

### Overview of genomic diversity in OB cattle

We annotated 15,722,811 biallelic SNPs and 1,580,878 Indels that were discovered in 49 OB cattle [17]. The average genome wide nucleotide diversity within the OB breed was 0.001637/bp. Among the detected variants, 546,419 (3.5%) SNPs and 307,847 (19.5%) Indels were found novel when compared to the 102,090,847 polymorphic sites of the NCBI bovine dbSNP database version 150.

Functional annotation of the polymorphic sites revealed that the vast majority of SNPs were located in either intergenic (73.8%) or intronic regions (25.2%). Only 1% of SNPs (160,707) were located in the exonic regions (Table 1). In protein-coding sequences, we detected 58,387, 47,249 and 1,264 synonymous, missense, and high impact SNPs, respectively. According to the SIFT scoring, 10,738 missense SNPs were classified as likely deleterious to protein function (SIFT score < 0.05). Among the high impact variants, we detected 580, 33, 106, 273 and 272 stop gain, stop lost, start lost, splice donor and splice acceptor variants, respectively. Deleterious and high impact variants were more frequent in the low than high allele frequency classes (Additional file 4).

**Table 1:**
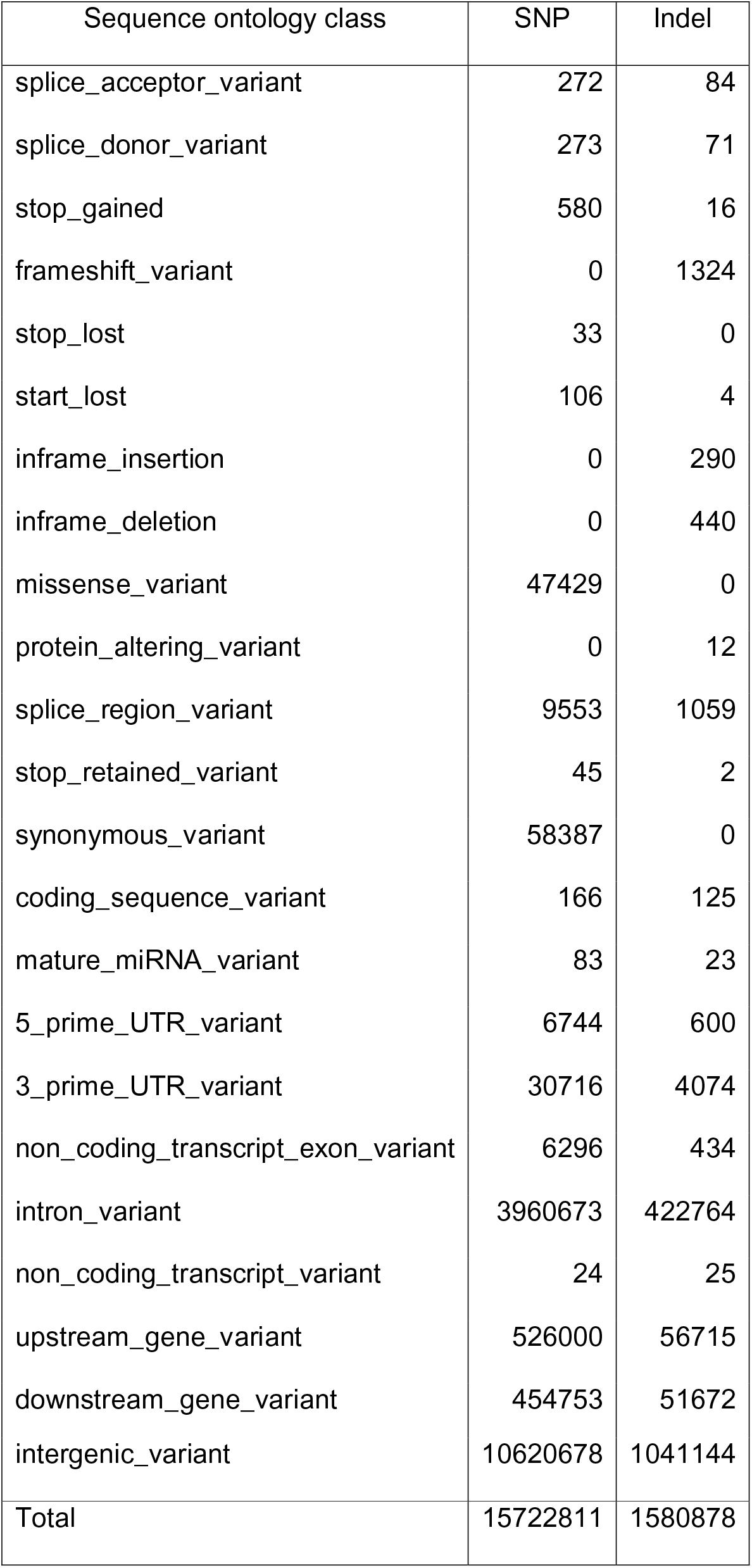
Number of SNPs and Indels in sequence ontology classes annotated using the VEP software Sequence ontology class SNP Indel.

| Sequence ontology class | SNP | Indel |
| --- | --- | --- |
| splice_acceptor_variant | 272 | 84 |
| splice_donor_variant | 273 | 71 |
| stop_gained | 580 | 16 |
| frameshift_variant | 0 | 1324 |
| stop_lost | 33 | 0 |
| start_lost | 106 | 4 |
| inframe_insertion | 0 | 290 |
| inframe_deletion | 0 | 440 |
| missense_variant | 47429 | 0 |
| protein_altering_variant | 0 | 12 |
| splice_region_variant | 9553 | 1059 |
| stop_retained_variant | 45 | 2 |
| synonymous_variant | 58387 | 0 |
| coding_sequence_variant | 166 | 125 |
| mature_miRNA_variant | 83 | 23 |
| 5_prime_UTR_variant | 6744 | 600 |
| 3_prime_UTR_variant | 30716 | 4074 |
| non_coding_transcript_exon_variant | 6296 | 434 |
| intron_variant | 3960673 | 422764 |
| non_coding_transcript_variant | 24 | 25 |
| upstream_gene_variant | 526000 | 56715 |
| downstream_gene_variant | 454753 | 51672 |
| intergenic_variant | 10620678 | 1041144 |
| Total | 15722811 | 1580878 |

The majority of 1,580,878 Indels were in either intergenic (72.7%) or intronic (26.7%) regions. Only 2,213 (0.14%) Indels affected coding sequences. Among these, 1,499 were classified as high impact variants including 1,324, 16, 4, 71 and 84 frameshift, stop gain, start lost, splice donor and splice acceptor variants, respectively. Similar to previous studies in cattle [14, 45], coding regions were enriched for Indels with lengths in multiples of three indicating that they are less likely to be deleterious to protein function than frameshift variants (Additional File 5).

### OMIA variants segregating in the OB population

We obtained genomic coordinates of 155 variants that are associated with Mendelian traits in cattle from the OMIA database to analyze if they segregate among the 49 OB cattle. It turned out that six OMIA variants were also detected in the 49 OB cattle including two variants in the *MOCOS* and *SLC45A2* genes that are associated with severe recessive disorders (Additional file 6). Two OB key ancestor bulls born in 1967 and 1974 (ENA SRA sample accession numbers SAMEA4827662 and SAMEA4827664) were heterozygous carriers of a single base pair deletion (BTA24:g.21222030delC) in the *MOCOS* gene (OMIA 001819-9913) that causes xanthinuria in the homozygous state in Tyrolean grey cattle [46]. Another two OB key ancestor bulls (sire and son; ENA SRA sample accession numbers SAMEA4827659 and SAMEA4827645) that were born in 1967 and 1973 were heterozygous carriers of two missense variants in *SLC45A2* (BTA20:g.39829806G>A and BTA20:g.39864148C>T) that are associated with oculocutaneous albinism (OMIA 001821-9913) in Braunvieh cattle [47].

### Runs of homozygosity and genomic inbreeding

Runs of homozygosity were analyzed in 33 OB animals that had an average sequencing depth greater than 10-fold. We found an average number of 2,044 ± 79 autosomal ROH per individual with an average length of 179 kb ± 17.6 kb. The length of the ROH ranged from 50 kb (minimum size considered, see methods) to 5,025,959 bp. On average, 14.58% of the genome (excluding sex chromosome) was in ROH (Additional file 7). Average genomic inbreeding for the 29 chromosomes ranged from 11.5% (BTA29) to 18.6% (BTA26) (Figure 1a).

**Figure 1.**
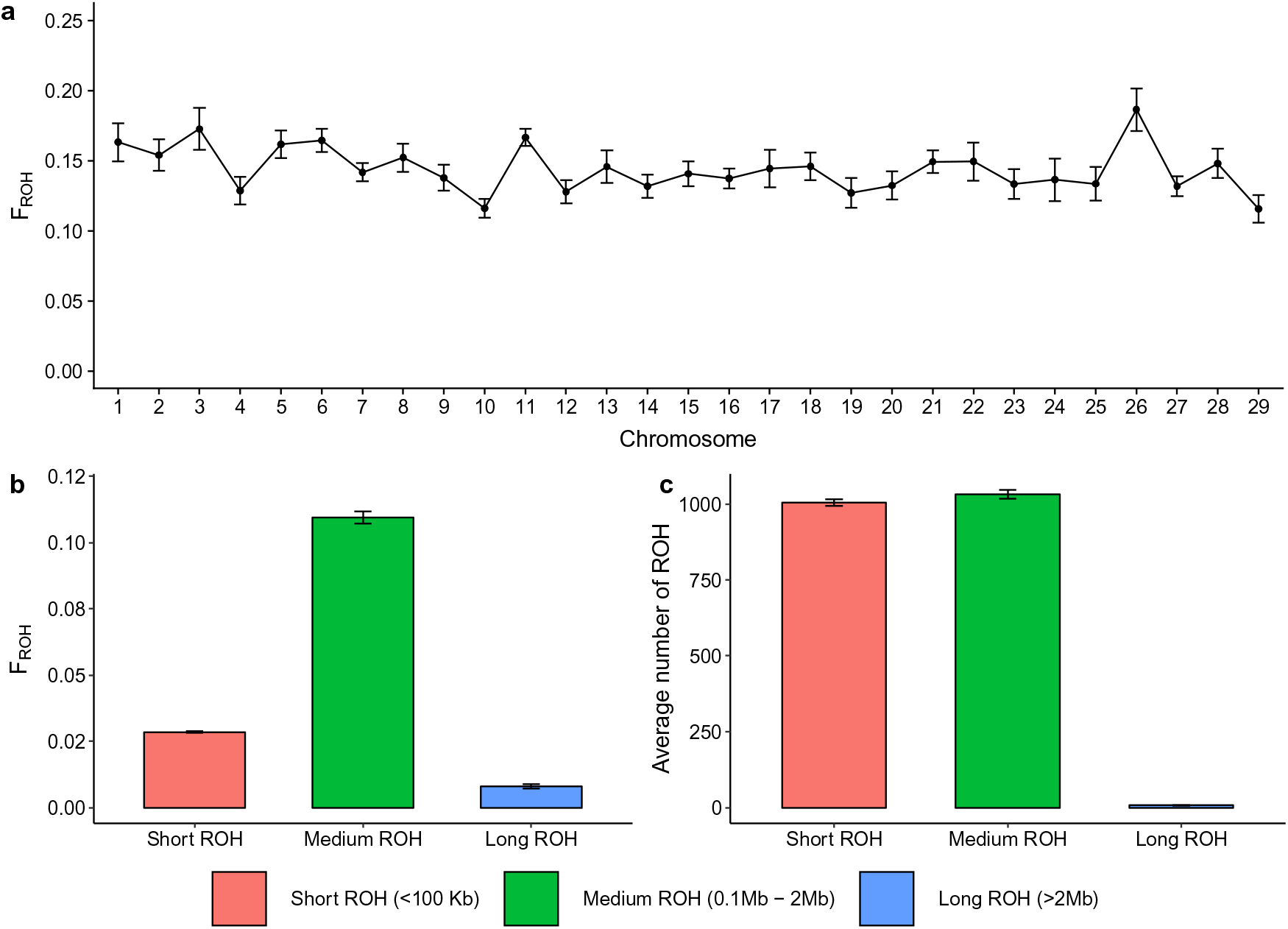
ROH in 33 OB cattle with average sequencing depth greater than 10-fold. (a) Average genomic inbreeding and corresponding standard error for the 29 autosomes. (b) Average genomic inbreeding (F_ROH_) calculated from short (50-100 kb), medium (0.1 - 2 Mb) and long (>2 Mb) ROH. (c) Average number of short, medium and long ROH. Genomic inbreeding (F_ROH_) was significantly (*P* =0.0002) higher in 20 animals born between 1990 and 2012 than in 13 animals born between 1965 and 1989 (0.16 vs. 0.14) (Additional file 8). The high F_ROH_ in animals born in more recent generations was mainly due to more long (>2 Mb; *P* =0.00004) and medium-sized ROH (0.1-1 Mb; *P* =0.001) (Figure 2).

**Figure 2.**
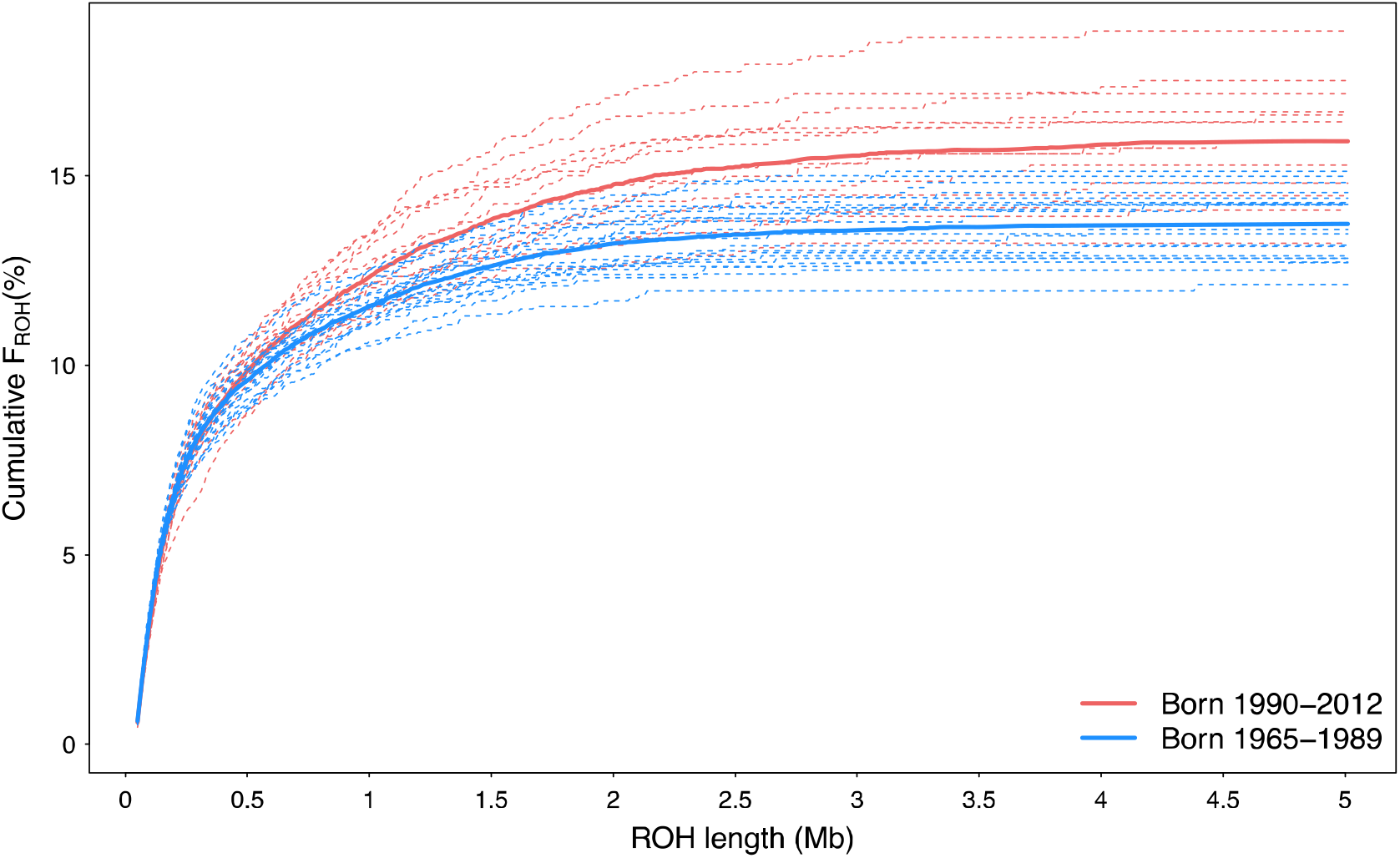
Cumulative genomic inbreeding (%) in animals born between 1965-1989 (blue lines) and 1990-2012 (red lines) from ROH sorted on length and binned in windows of 10 kb. Thin dashed lines represent individuals and thick solid lines represent the average cumulative genomic inbreeding of the two groups of animals.

In order to study the demography of the OB population, we calculated the contributions of short, medium and long ROH to the total genomic inbreeding (Additional file 7). The medium-sized ROH were the most frequent class (50.46%), and contributed most (75.01%) to the total genomic inbreeding. While short ROH occurred almost as frequent (49.17%) as medium-sized ROH, they contributed only 19.52% to total genomic inbreeding (Figure 1b&c; Additional file 7). Long ROH were rarely (0.36%) observed among the OB key ancestors and contributed little (5.47%) to total genomic inbreeding. The number of long ROH was correlated (r=0.77) with genomic inbreeding.

### Signatures of selection

We identified candidate signatures of selection using two complementary methods: the composite likelihood ratio (CLR) test and the integrated haplotype score (iHS) (Figure 3a & 3b). The CLR test detects ‘hard sweeps’ where genomic regions encompass beneficial adaptive alleles that recently reached fixation [48]. The iHS detects ‘soft sweeps’ at genomic regions where selection for beneficial alleles is still ongoing [49, 50]. We detected 95 and 162 candidate regions of signatures of selection (*P* < 0.005) using CLR and iHS, respectively, encompassing 12.56 Mb and 12.48 Mb (Additional file 9; Additional file 10). These candidate signatures of selection were not evenly distributed over the genome (Figure 3c). Functional annotation revealed that 136 and 157 protein-coding genes overlapped with 50 and 86 candidate regions from CLR and iHS analyses, respectively. All other candidate signatures of selection were located in intergenic regions. Closer inspection of the top selection regions of both analyses revealed that 16 CLR candidate regions overlapped with 25 iHS candidate regions on chromosomes 5, 7, 11, 14, 15, 17 and 26 (Figure 3c) encompassing 35 coding genes (Additional File 11).

**Figure 3.**
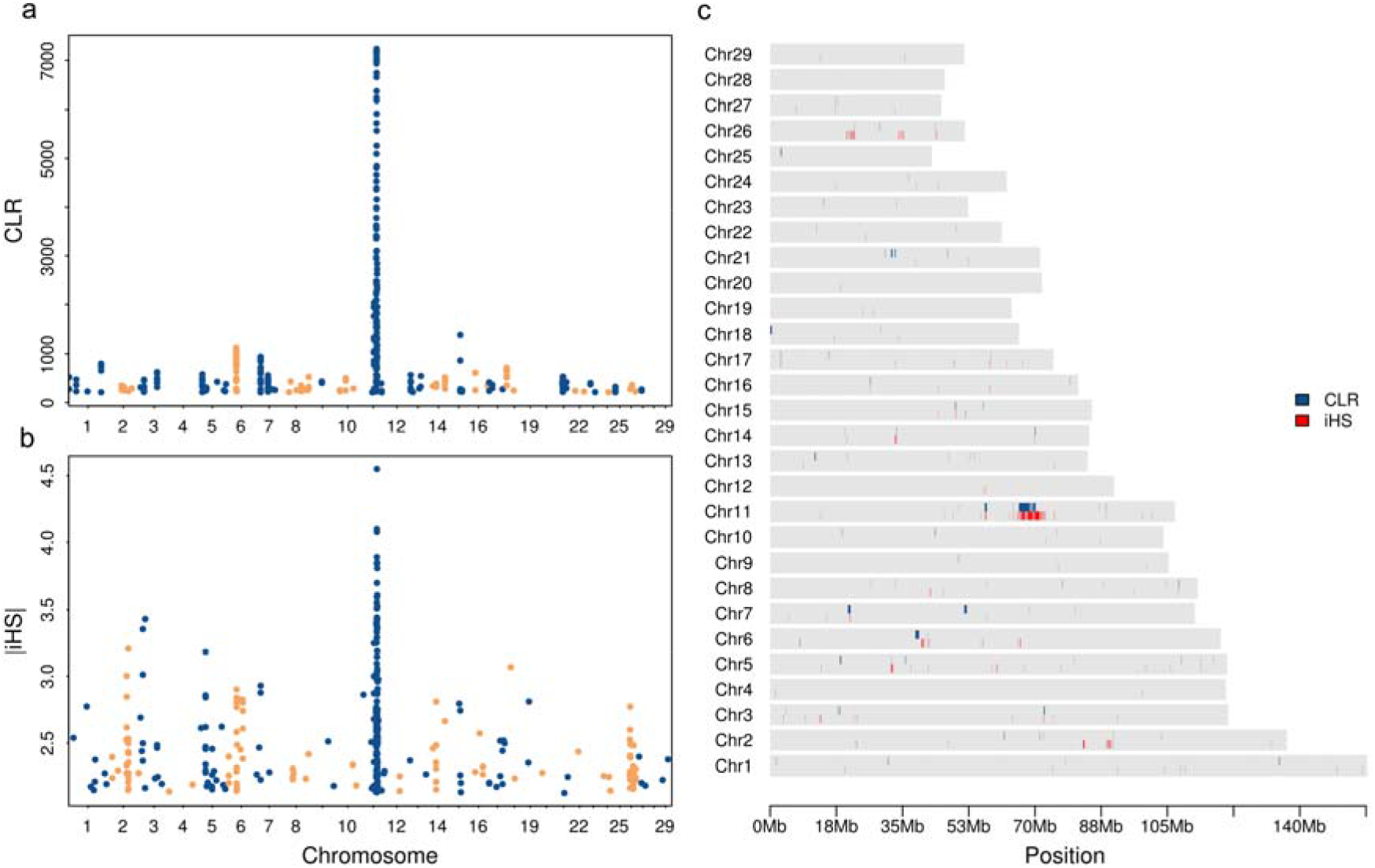
Genome wide distribution of top 0.5% signatures of selection from CLR (a) and iHS (b) analyses and their overlap (c). Each point represents a non-overlapping window of 40 kb along the autosomes.

### Top candidate signatures of selection

On chromosome 11, we identified 12 and 36 candidate regions of selection using CLR and iHS analyses, respectively. The top CLR candidate region (*P_CLR_* = 3.1 x 10^-5^) was located on chromosome 11 between 66 Mb and 68.5 Mb (Figure 4a) and it encompassed 24 protein-coding genes (Additional file 9). The same region was also in ROH in 77% of 33 animals that were sequenced at high coverage. The peak of this top CLR region was located between 67.5 and 68.2 Mb and it contained several adjacent windows with CLR values higher than 5,000 (*P_CLR_* < 0.003). The top region encompassed 5 genes (Figure 4a & 4e). The variant density in the top region was low and SNP allele frequency was skewed which is typical for the presence of a hard sweep (Figure 4c). The top iHS candidate region was located on chromosome 11 between 68.4 and 69.2 Mb (*P_iHS_* = 3.2 x 10^-5^) encompassing 7 genes (Figure 4b & 4f). The allele frequencies of the SNPs within the top iHS region are approaching fixation indicating ongoing selection possibly due to hitchhiking with the neighboring hard sweep (Figure 4d).

**Figure 4.**
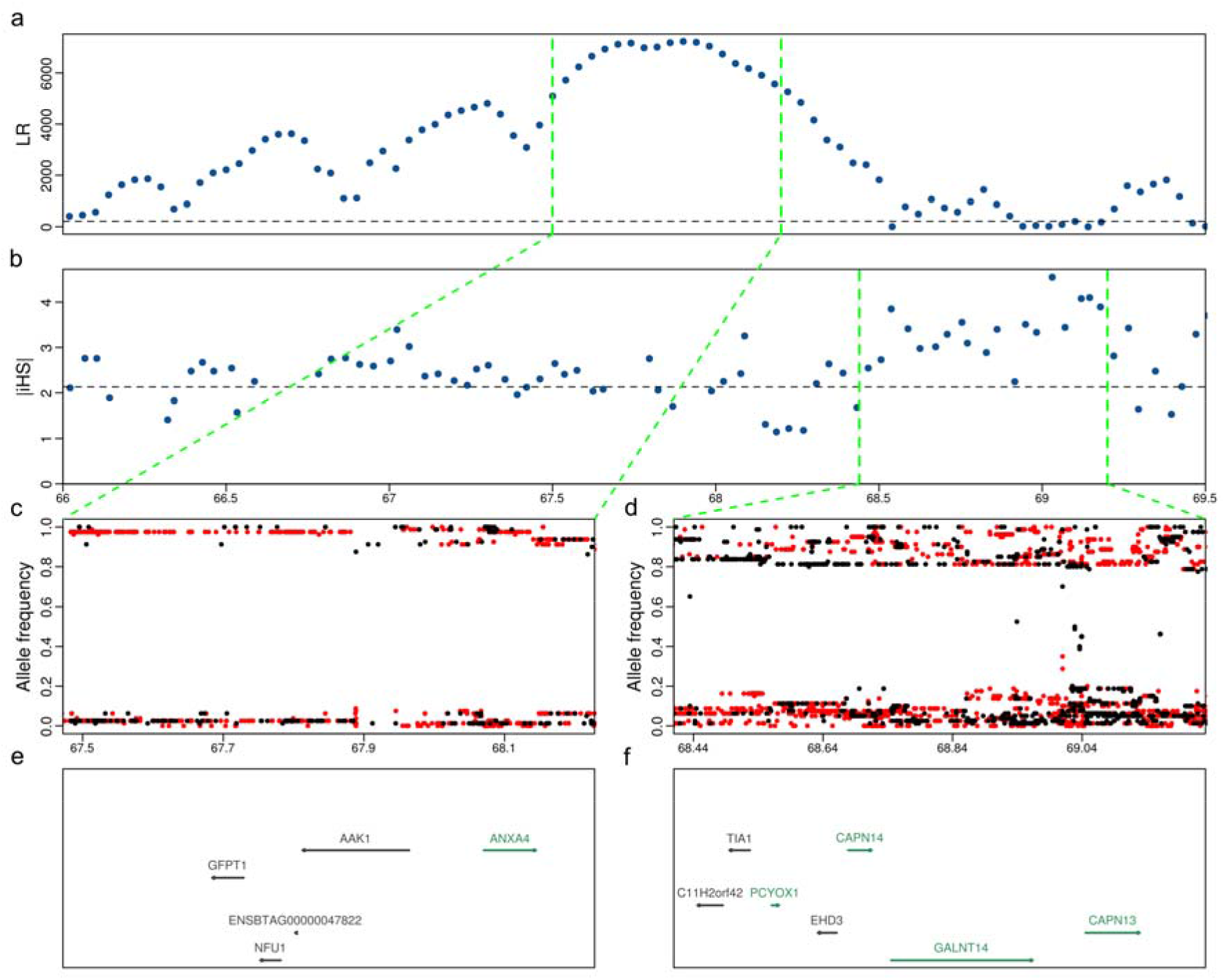
Detailed view of a top candidate selection region on chromosome 11 in OB that was detected using CLR tests (a) and iHS (b). Each point represents a non-overlapping window of 40 kb. The dotted horizontal lines indicate the cutoff values (top 0.5%) for CLR (210) and iHS (2.13) statistics. The allele frequencies of the derived (red) or alternate alleles (black) (c and d) and genes (e and f) in the peak region (67.5-68.2 Mb) of the top CLR (66 to 68.5 Mb) and iHS (68.4-69.2 Mb) regions. Green and black colour indicates genes on the forward and reverse strand of DNA, respectively.

Another striking CLR signal (*P_CLR_* = 0.0012) was detected on chromosome 6 between 38.5 and 39.4 Mb. This genomic region encompasses the *DCAF16, FAM184B, LAP3, LCORL, MED28* and *NCAPG* genes, and the window with the highest CLR value overlapped the *NCAPG* gene (Figure 5a & 5c). This signature of selection coincides with a QTL that is associated with stature, feed efficiency and fetal growth [51–53]. Most SNPs detected within this region were fixed for the alternate allele in the OB key ancestor animals of our study. For instance, all 49 sequenced OB cattle were homozygous for the Chr6:38777311 G-allele which results in a likely deleterious (SIFT score 0.01) amino acid substitution (p.I442M) in the *NCAPG* gene that is associated with increased pre- and postnatal growth and calving difficulties [51].

**Figure 5.**
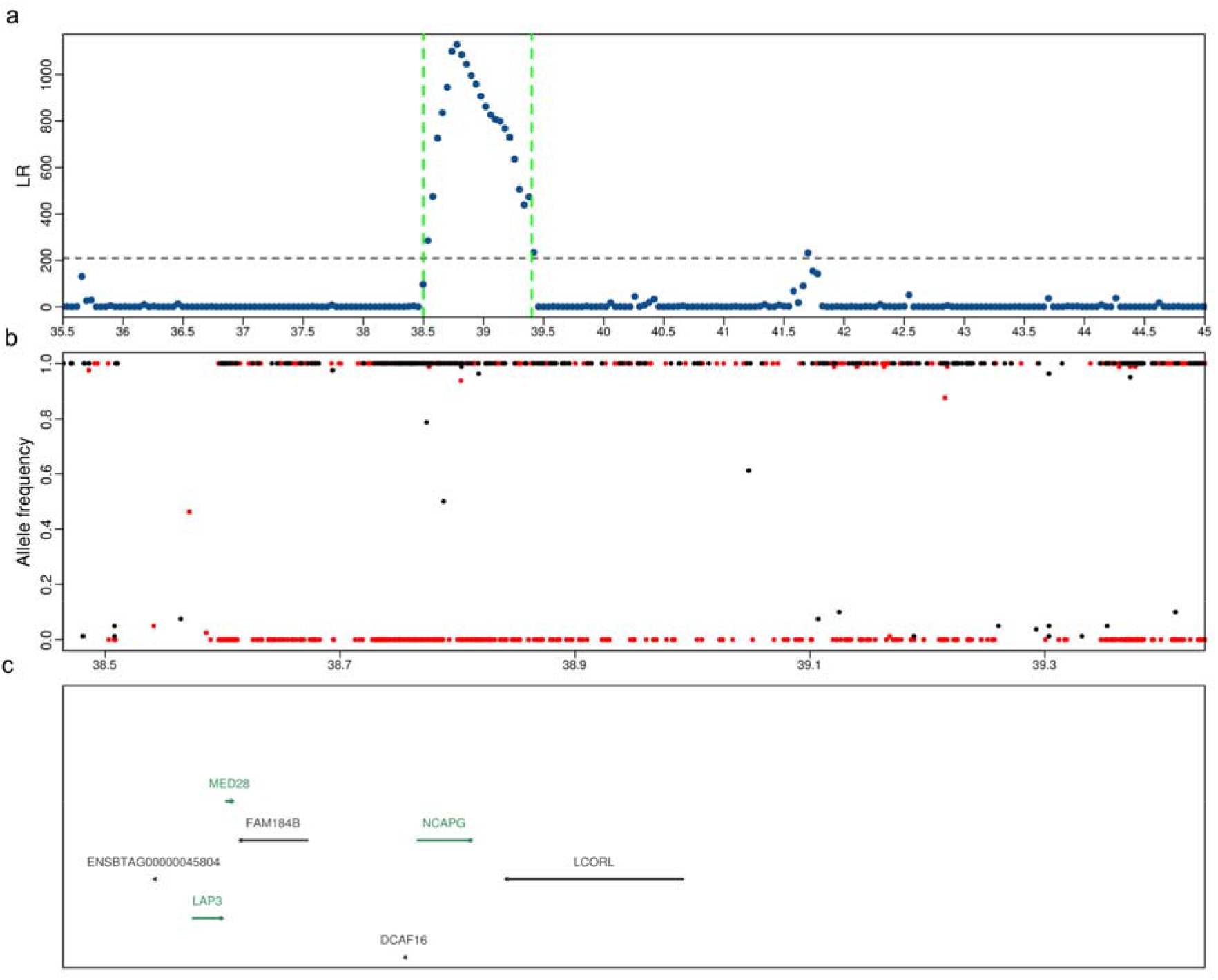
Top CLR candidate region on chromosome 6 (a). Each point represents a non-overlapping window of 40 kb. The frequencies of the derived (red) or alternate alleles (black) (c) and genes (e) annotated between 38.5 and 39.4 Mb. Green and black colour indicates genes on the forward and reverse strand of DNA, respectively.

### GO enrichment analysis

Genes within candidate signatures of selection from CLR and iHS analyses were enriched (after correcting for multiple testing) in the panther pathway (P00011) related to “Blood coagulation”. Genes within candidate signatures of selection from CLR tests were also enriched in the pathway “P53 pathway feedback loops 1” (Additional file 12). Although we did not find any enrichment of GO-slim biological processes after correcting for multiple testing, 21 GO-slim biological processes including cellular catabolic processes, oxygen transport and different splicing pathways were nominally enriched for genes within CLR candidate signatures of selection and 14 GO-slim biological processes including nervous system, sensory perception (olfactory receptors) and multicellular processes were nominally enriched for genes within iHS candidate signatures of selection (Additional file 12).

### QTL enrichment analysis

We investigated if candidate selection regions overlapped with trait-associated genomic regions using QTL information curated at the Animal QTL Database (Animal QTLdb). We found that 74.7% and 83.9% of CLR and iHS candidate signatures of selection, respectively, were overlapping at least one QTL (Additional file 13). We tested for enrichment of these signatures of selection in QTL for six trait classes: exterior, health, milk, meat, production, and reproduction using permutation. It turned out that QTL associated with meat quality (*P_CLR_* = 0.0004, *P_iHS_* = 0.0003) and production traits (*P_CLR_* = 0.0027, *P_iHS_* = 0.0039) were significantly enriched in both CLR and iHS candidate signatures of selection. We did not detect any enrichment of QTL associated with milk, reproduction, health, and exterior traits neither in CLR nor in iHS candidate signatures of selection.

## Discussion

We discovered 107,291 variants in coding sequences of 49 sequenced OB cattle. In agreement with previous studies in cattle [14, 54], missense deleterious and high impact variants occurred predominantly at low allele frequency likely indicating that variants which disrupt physiological protein functions are removed from the population through purifying selection [55]. However, deleterious variants may reach high frequency in livestock populations due to the frequent use of individual carrier animals in artificial insemination [56], hitchhiking with favorable alleles under artificial selection [57, 58], or demography effects such as population bottlenecks [59]. Because we predicted functional consequences of missense variants using computational inference, they have to be treated with caution in the absence of experimental validation [60]. High impact variants that segregated among the 49 sequenced OB key ancestors were also listed as Mendelian trait-associated variants in the OMIA database. For instance, we detected frameshift and missense variants in *MOCOS* and SLC45A2 that are associated with recessive xanthinuria [46] and oculocutaneous albinism [47], respectively. To the best of our knowledge, calves neither with xanthinuria nor oculocutaneous albinism have been reported in the Swiss OB cattle population. The absence of affected calves is likely due to the low frequencies of the deleterious alleles and avoidance of matings between closely related heterozygous carriers. Among 49 sequenced cattle, we detected only two bulls that carried the disease-associated *MOCOS* and *SLC45A2* alleles in the heterozygous state. However, the frequent use of individual carrier bulls in artificial insemination might result in an accumulation of diseased animals within short time even when the frequency of the deleterious allele is low in the population [61]. Because the deleterious alleles were detected in sequenced key ancestor animals that were born decades ago, we cannot preclude that they were lost due to genetic drift or during the recent population bottleneck in OB (Additional file1). A frameshift variant in *SLC2A2* (NM_001103222:c.771_778delTTGAAAAGinsCATC, rs379675307, OMIA 000366-9913) causes a recessive disorder in cattle that resembles human Fanconi-Bickel syndrome [62–64]. Recently, the disease-causing allele was detected in the homozygous state in an OB calf with retarded growth due to liver and kidney disease [65]. We did not detect the disease-associated allele in our study. This may be because it is located on a rare haplotype that does not segregate in the 49 sequenced cattle. Most of the sequenced animals of the present study were selected for sequencing using the key ancestor approach, as their genes contributed significantly to the current population [17, 66]. More sophisticated methods to select animals for sequencing might prioritize rare haplotypes, thus increasing the likelihood to detect rarer alleles when the sequencing budget is constrained [67–69].

### Genomic diversity and genomic inbreeding

Original Braunvieh is a local cattle breed with approximately 10,000 cows registered in the breeding population and 4,500 calves entering the herd book every year (Additional file 1). In spite of the small population size, the nucleotide diversity (π = 1.6 × 10^-3^) is higher in OB than many taurine cattle breeds with considerably more breeding animals including Holstein, Jersey and Fleckvieh (∼1.2-1.4 × 10) [70, 71]. However, nucleotide diversity is lower in OB than African indigenous cattle breeds (2.0-4.0 × 10^-3^), New Danish Red (1.7 × 10^-3^) and Yakutian cattle breeds (1.7 × 10^-3^) [71–73]. The average F_ROH_ estimated from WGS data was 0.14 in OB. This is lower than WGS-based F_ROH_ in Holstein (0.18), Jersey (0.24), Old Danish Red (0.23) and Belgian Blue (0.3) cattle [74, 75]. However, the genomic inbreeding is slightly higher in OB than New Red Danish cattle (0.11), an admixed breed that contains genes from old Danish and other red breeds [74]. The relatively high genomic diversity of OB cattle is assumed to be the result of many different sires contributing to the gene pool due to frequent use of natural mating [10]. Our WGS based estimate of F_ROH_ (0.14) is substantially higher than previous estimates obtained using 50K SNP microarray data (F_ROH_= 0.029, [10]) for the same population. Genotype data obtained using SNP microarrays with medium density (e.g., BovineSNP50) facilitate to detect long ROH (>1 Mb). However, due to low SNP density (∼1 SNP per 50 kb) detecting short ROH is not possible using microarray-derived genotype data. In our data, short and medium-sized ROH accounted for 80.48% of total inbreeding. Most short and medium-sized ROH are not reliably detectable with the SNP microarrays that were used to quantify F_ROH_ in Signer-Hasler et al. [10], resulting in an underestimation of genomic inbreeding. Our estimate of the genomic inbreeding using WGS variants also includes short and medium-sized ROH that were previously missed using SNP array data, thus representing a realistic estimate of total genomic inbreeding in OB cattle.

Apart from genomic inbreeding, ROH also provide information about population and individual demography based on length and number of ROH [76–78]. Our findings show that medium-sized ROH that reflect historical inbreeding contribute most to the genomic inbreeding of the current OB population. The minor contribution of long ROH to the genomic inbreeding indicates that recent inbreeding is relatively low in OB possibly due to use of many sires in natural matings as suggested by Hagger [9] and Signer-Hasler et al. [10]. Our results based on ROH inferred from WGS variants corroborate that genomic inbreeding is lower in OB than most mainstream breeds [10]. However, comparing the number and size distribution of ROH across studies is subject to bias because misplaced genomic segments might break ROH into multiple small- and medium-sized ROHs and different ROH-detection approaches yield results that are not readily comparable [21, 27, 76, 79, 80]. Genomic inbreeding is increasing in the OB population in recent years mainly due to an increase in occurrence of long ROH. The recent population bottleneck in the OB population (Additional file 1) might promote matings between closely related animals that caused inbreeding to increase in recent generations. In this regard, genome-based mating strategies seem to be warranted to achieve sufficient genetic gain while maintaining genetic diversity and avoiding matings between carriers of disease-associated alleles [81, 82].

### Signatures of selection

With WGS data, we were able to identify more signatures of selection compared to SNP array data [10, 83], even though we used only 9 million SNP for which we could readily assign ancestral and derived alleles [33]. Using two complementary approaches, we found several new and known candidate regions that seem to be targets of recent or ongoing selection in OB. Many signatures of selection were located in non-coding regions corroborating that selection frequently acts on regulatory sites [16]. However, it is possible that an improved annotation of the bovine genome might place these regions in yet to be annotated coding regions. We applied methods to detect signatures of selection that depend on frequency changes of alleles (CLR) and haplotypes (iHS). The detected signatures of selection may be confounded by other evolutionary forces including genetic drift and background selection [84–86].

We detected candidate selection regions in OB cattle that harbor genes associated with stature or milk production (*NCAPG, LCORL, LAP3*), feed efficiency or lipid metabolism (*R3HDM1, AOX1*), and unknown functions (*SLC25A33, TMEM201*) that were previously reported to be targets of selection in taurine and indicine cattle breeds [16, 87–89]. The presence of signatures of selection that are common in several breeds indicates that selection at these regions has happened either before the breeds diverged or independently after the formation of breeds [16, 89]. A number of genes that are targets of selection in various cattle breeds are associated with either coat colour (*MC1R*, *KIT*), milk production (*DGAT1*, *ABCG2*, *GHR*) or stature (*PLAG1*) [40, 71, 89, 90]. These genes were not detected within the top 0.5% CLR and iHS windows in OB cattle possibly either due to absence of trait-associated genomic variation in our data or these genes are not under selection in OB cattle. While some cattle breeds including Holstein and Fleckvieh are selected for particular coat colour patterns [40, 91], animals with variation in coat colour are rarely observed in the OB cattle breed [92]. Moreover, due to the use of OB cattle for both milk and beef production under extensive conditions, the milk production-associated variants that are under strong artificial selection in many dairy breeds seem to be less important in OB cattle [88].

Some of the genes (*PLAG1*, *DGAT1*, *ABCG2*, *GHR*) that have been reported to be targets of selection in specialized breeds contain well-known variants that contribute to the genetic variation of economically important traits. We investigated if these variants segregate in our data although they were not detected in our selection signature analysis. A number of variants in high linkage disequilibrium stimulate the expression of *PLAG1*, thus increasing pre- and postnatal growth in cattle [52, 53, 93]. Among 14 candidate causal variants for the *PLAG1* QTL, six were fixed for the stature-decreasing alleles in our study (Additional File 14). The other candidate causal variants were either fixed for the stature-increasing allele or segregated at low allele frequency. This pattern indicates that a recombinant haplotype might segregate in Swiss OB cattle that could facilitate fine-mapping of this region. Among known mutations affecting milk production traits, a mutation in *ABCG2* (p.Y581S, rs43702337 at 38,027,010 bp) [94] did not segregate in our population of Swiss OB cattle which corroborates previous findings in Brown Swiss cattle [95]. A variant (BTA14, g.1802265_1802266GC>AA, p.A232K) of the *DGAT1* gene is associated with milk production traits in cattle [96, 97]. The milk fat-enhancing and milk yield-lowering lysine-allele segregates in OB cattle at low frequency (0.03). A missense variant (BTA20, g.31909478A>T, p.Y279F) in the *GHR* gene is associated with milk protein percentage [98]. The protein fat percentage-lowering T-allele segregates at low frequency (0.06) in OB cattle.

We observed a striking signature of selection on chromosome 11 that has previously been detected in the Swiss Fleckvieh, Simmental, Eringer and Evolèner breeds using microarray-called genotypes [10, 83]. Our results in OB cattle indicate that this region harbors a rapid sweep which seems to act on alleles with selection advantage [49]. While large sweeps are easy to detect using dense sequencing data, pinpointing causal alleles underpinning such regions remains challenging. Most of the variants in such regions are either fixed or segregate at very low frequency [16] which we also observed for the signature of selection on chromosome 11. In our study, the signature of selection on chromosome 11 encompassed millions of nucleotides (between 66 and 72 Mb) and many genes, rendering the identification of underpinning genes and variants a difficult task. The windows with the highest CLR and |iHS| values did not encompass *PROKR1*, which was previously suggested to be the target of selection at this region due to its association with fertility [10, 83]. However, closer inspection of the sequence variants detected in our study revealed that a stop-gained variant in *PROKR1* (g.66998234C>A, rs476744845, p.Y293*) segregates at high frequency in the 49 sequenced OB key ancestors. Yet, it remains to be elucidated, if the presence of a high-impact variant in immediate proximity to a massive selective sweep is causal or just due to hitchhiking. The window with the largest |iHS| value on chromosome 11 was right next to the *CAPN13* gene which is associated with meat tenderness and was also suggested as a potential target of selection by Signer-Hasler et al. [10].

Genes within the top 0.5% CLR and iHS windows were enriched in pathways related to reactive oxygen species, metabolic process, blood coagulation and nervous system, indicating that the identified regions under selection might harbor genomic variants that confer adaptive advantage to harsh environments. Moreover, QTL associated with meat quality and production traits including feed efficiency and body weight were enriched in selection signatures possibly indicating that OB cattle harbor variants that enabled them to adapt to particular feed conditions. Combining results from selection signature and association analyses might reveal phenotypic characteristics associated with genomic regions that showed evidence of past or ongoing selection [40], thus providing additional hints why particular genomic regions are under selection in OB cattle.

## Conclusions

We provide a comprehensive overview of genomic variation segregating in the Swiss OB cattle population using sequencing data of 49 key ancestor bulls. In spite of the small population size, genetic diversity is higher and genomic inbreeding is lower in OB than many other mainstream cattle breeds. However, genomic inbreeding is increasing in recent generations mainly due to large ROH which should be considered in future management of this breed. Finally, this study highlights regions that show evidence of past and ongoing selection in OB which are enriched for QTL related to meat quality and production traits and pathways related to blood coagulation, cellular metabolic process, and nervous system.

## List of abbreviations

WGS: Whole-genome sequencing
CLR: Composite likelihood ratio
iHS: Integrated haplotype score
OB: Original Braunvieh
OMIA: Online Mendelian Inheritance in Animals
QTL: Quantitative trait loci
ROH: runs of homozygosity
SNP: Single nucleotide polymorphism

## Declarations

### Ethics approval and consent to participate

Not applicable

### Consent for publication

Not applicable

### Availability of data and materials

Raw sequencing read data of all animals are available from the European Nucleotide Archive (ENA) (http://www.ebi.ac.uk/ena) under primary accession PRJEB28191 (SAMEA4827645-SAMEA4827674 and SAMEA5059741-SAMEA5059759).

### Funding

This study was financially supported by the Federal Office for Agriculture (FOAG), Bern. The funding body was not involved in the design of the study and collection, analysis, and interpretation of data and in writing the manuscript.

### Author’s contributions

Analysis of whole-genome sequencing data: MB NKK CD HP; Conceived and designed the experiments: MB NKK HP; Wrote the paper: MB HP; Read and approved the final version of the manuscript: all authors

## Acknowledgements

We thank Braunvieh Schweiz for providing pedigree and genotype data of Original Braunvieh cattle.

## Supplementary Information

**Additional file 1 File S1**

File format: .docx

Title: Original Braunvieh herdbook population

Description: Number of female calves entering the OB herdbook between 1980 and 2016.

**Additional file 2 File S2**

File format: .pdf

Title: Runs of homozygosity in 49 OB cattle

Description: a) Total genome fraction in ROH in 49 cattle with high (<10x) and low (<10x) coverage (b) Phred confidence score for ROH in 33 cattle sequenced at average sequencing depth higher than 10-fold. Red dots indicate mean confidence scores for ROH.

**Additional file 3 File S3**

File format: .xlsx

Title: Bovine QTL information downloaded from the AnimalQTL database. Description: Number of QTL for each trait. Traits are grouped in six trait categories.

**Additional file 4 File S4**

File format: .pdf

Title: Allele frequency distribution in different functional annotations

Description: Allele frequency of SNPs with different consequences according to VEP prediction, like high impact, deleterious (missense SNP with SIFT score < 0.05). tolerated (missense SNPs with SIFT score > 0.05) and synonymous SNPs.

**Additional file 5 File S5**

File format: .pdf

Title: Distribution of length of Indels

Description: a) Number of Indels (x1000) with size less than 12 bp detected according to the number of affected bases. b) number of Indels detected in coding sequences.

**Additional file 6 File S6**

File format: .xlsx

Title: OMIA variants detected in OB

Description: Six OMIA variants detected in 49 sequenced OB animals with their respective frequency and information.

**Additional file 7 File S7**

File format: .xlsx

Title: ROH statistics of each animal (high coverage)

Description: Number, length, average length and genomic fraction of all ROH in each animal. Also categorized in short, medium and long ROH category.

**Additional file 8 File S8**

File format: .pdf

Title: Genomic inbreeding in OB cattle stratified by birth year

Description: Genomic inbreeding in two groups of animals born either between 1960 and 1989 or between 1990 and 2012.

**Additional file 9 File S9**

File format: .xlsx

Title: Candidate selection signatures detected using CLR

Description: Genomic coordinates, CLR values, p values and encompassed genes for 95 candidate selection signatures.

**Additional file 10 File S10**

File format: .xlsx

Title: Candidate selection signatures detected using iHS

Description: Genomic coordinates, |iHS| values, p values and encompassed genes for 162 candidate selection signatures.

**Additional file 11 File S11**

File format: .xlsx

Title: Overlap between the top CLR and iHS selection signatures

Description: Overlapped regions and genes between CLR and iHS candidate selection regions

**Additional file 12 File S12**

File format: .xlsx

Title: Summary of Gene Ontology enrichment analysis

Description: PANTHER and GO-Slim pathways enriched using genes encompassed in signatures of selection from CLR and iHS analyses.

**Additional file 13 File S13**

File format: .xlsx

Title: Overlap between QTL and signatures of selection

Description: QTL (from all 6 categories) that overlapped with CLR and iHS selection signatures.

**Additional file 14 File S14**

File format: .xlsx

Title: Frequency of candidate causal variants for a stature QTL on BTA14 Description: Genomic coordinates and allele frequencies of 14 variants nearby bovine *PLAG1* that were reported as candidate causal variants for a stature QTL in cattle by Karim et al. [95].

