## Supplementary figures and images for "Assessing genomic diversity and signatures of selection in Original Braunvieh cattle using whole-genome sequencing data"

### File_S1

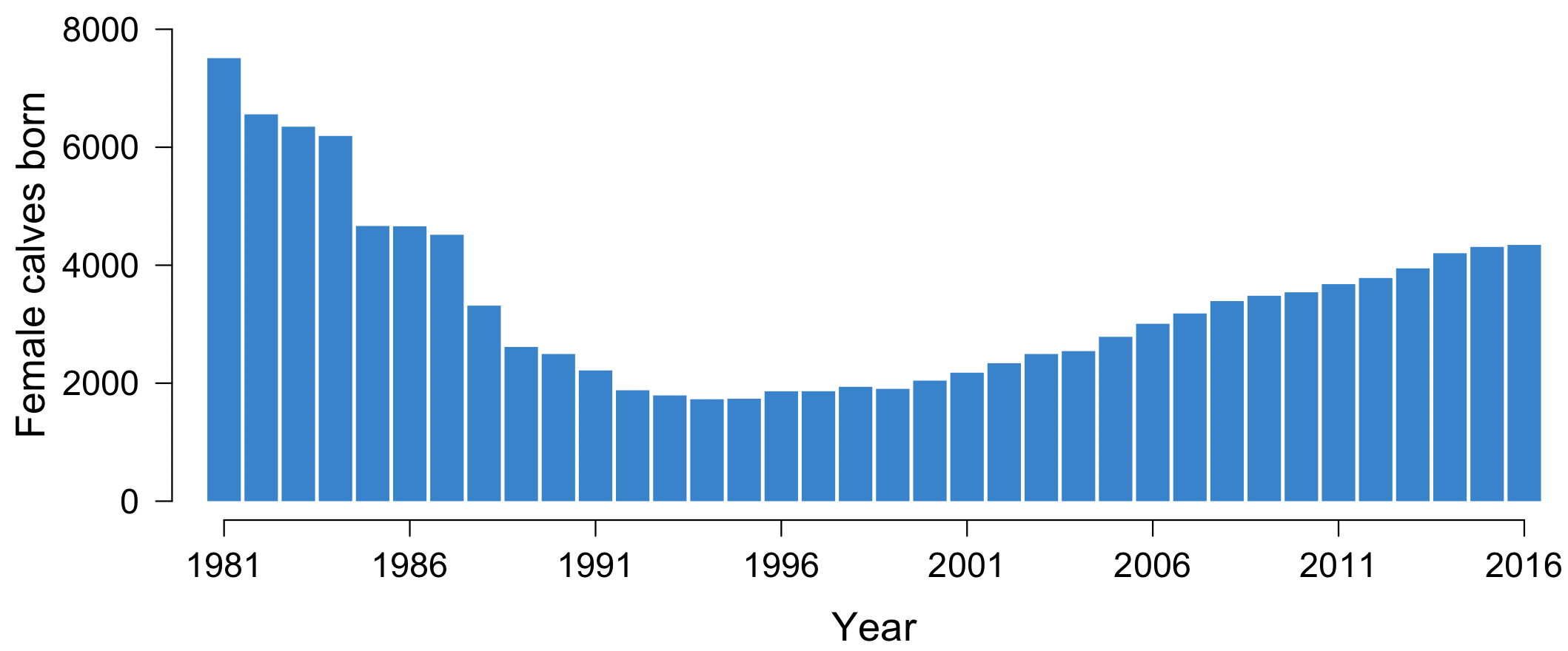

### File_S2

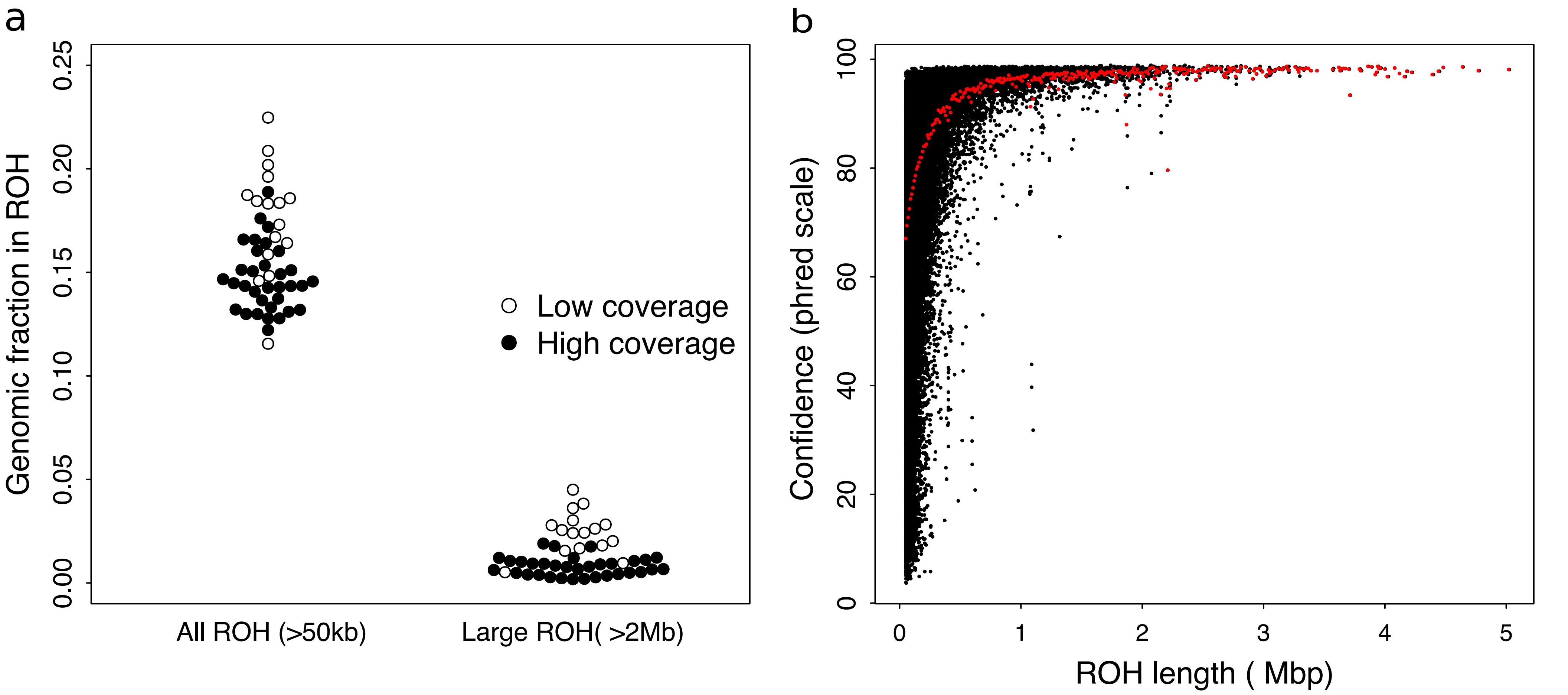

### File_S4

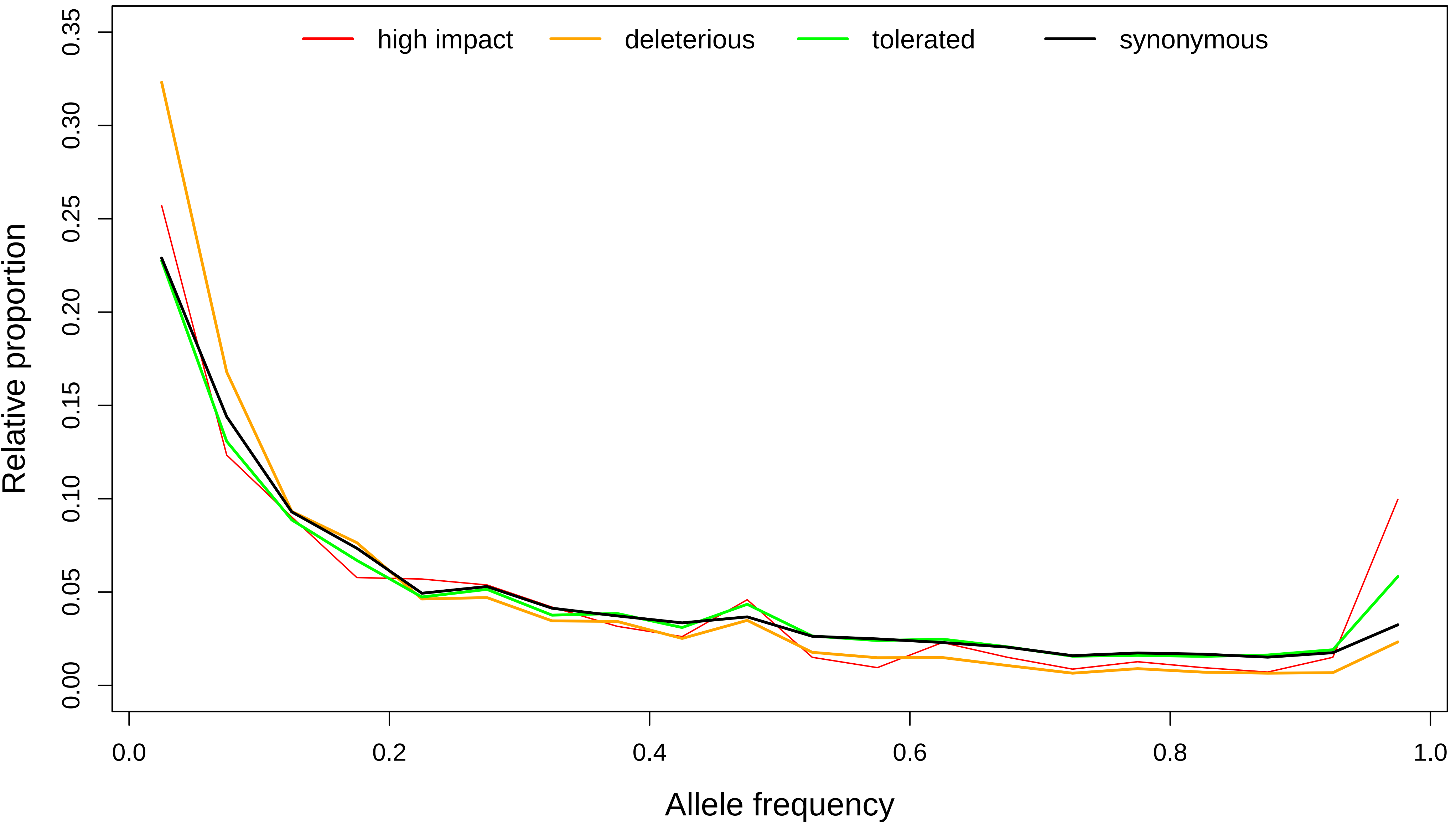

### File_S5

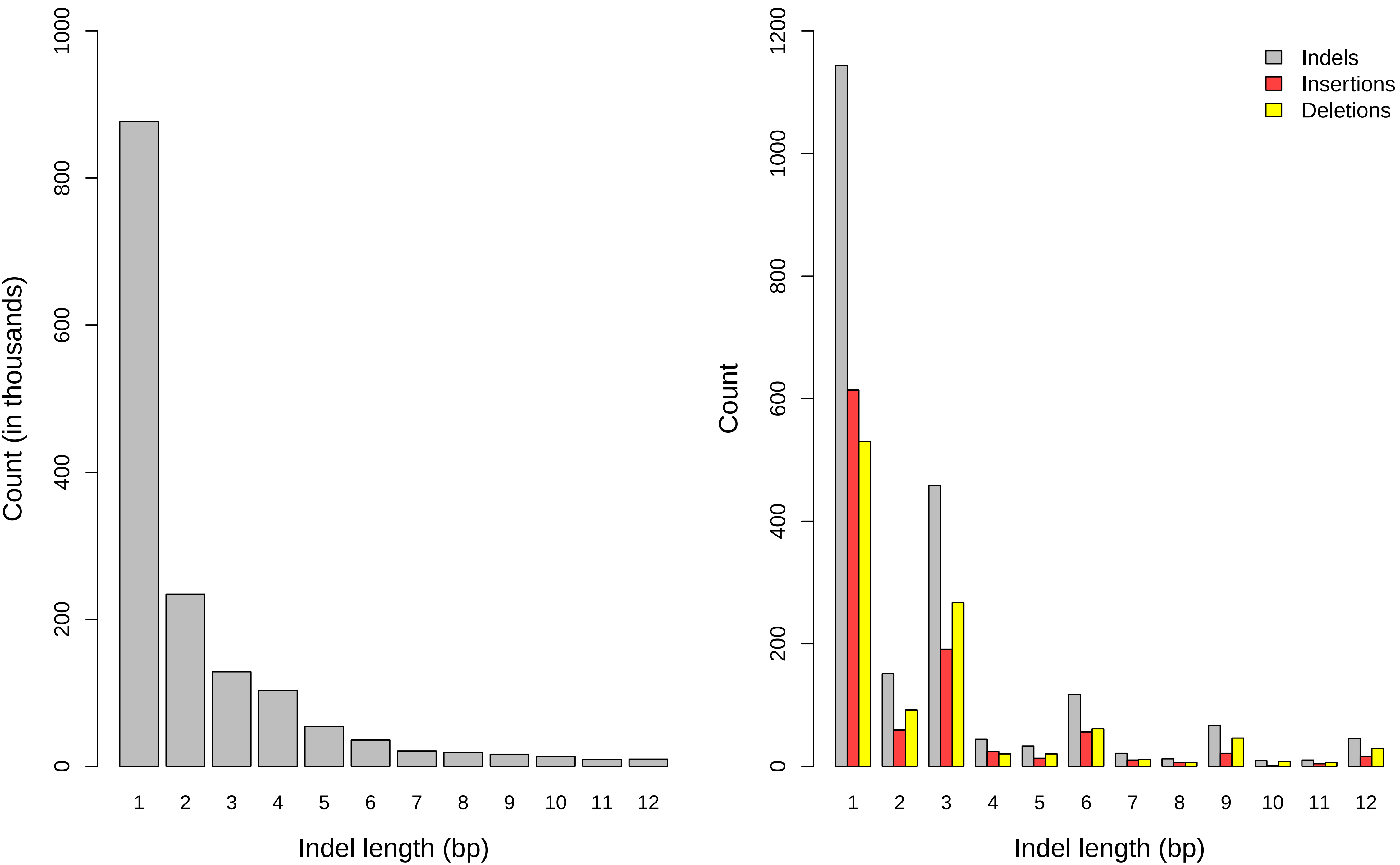

### File_S8

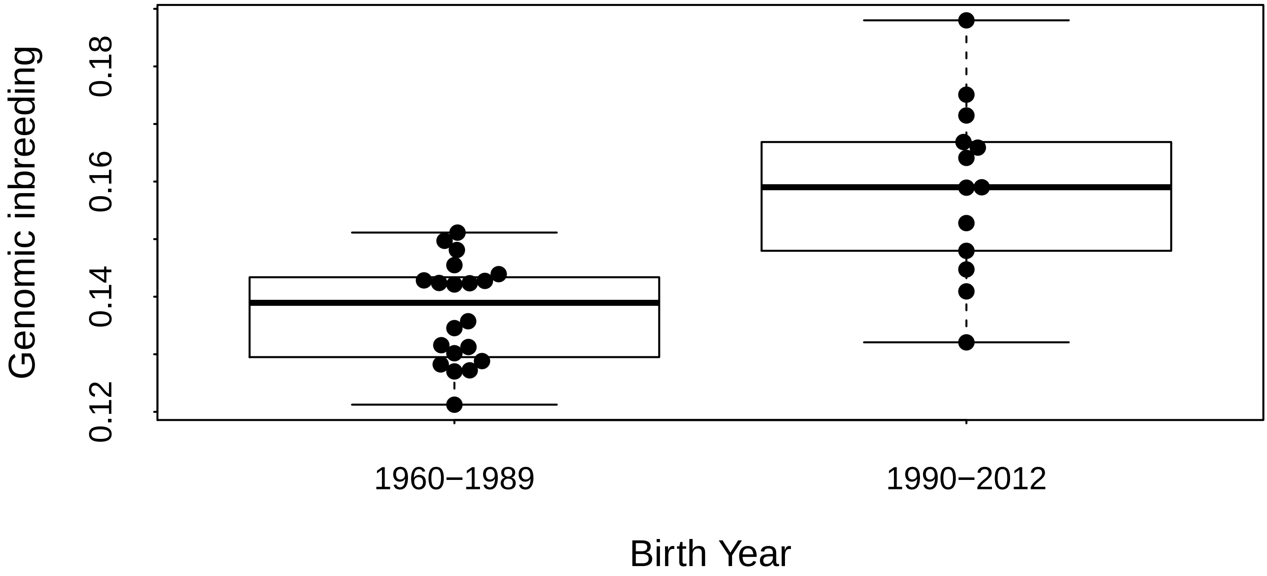
